# Is it reasonable to account for population structure in genome-wide association studies?

**DOI:** 10.1101/647768

**Authors:** Bongsong Kim

## Abstract

Population structure is widely perceived as a noise factor that undermines the quality of association between an SNP variable and a phenotypic variable in genome-wide association studies (GWAS). The linear model for GWAS generally accounts for population-structure variables to obtain the adjusted phenotype which has less noise. Its result is known to amplify the contrast between significant SNPs and insignificant SNPs in a resultant Manhattan plot. In fact, however, conventional GWAS practice often implements the linear model in an unusual way in that the population-structure variables are incorporated into the linear model in the form of continuous variables rather than factor variables. If the coefficients for population-structure variables change across all SNPs, then each SNP variable will be regressed against a differently adjusted phenotypic variable, making the GWAS process unreliable. Focusing on this concern, this study investigated whether accounting for population-structure variables in the linear model for GWAS can assure the adjusted phenotypes to be consistent across all SNPs. The result showed that the adjusted phenotypes resulting across all SNPs were not consistent, which is alarming considering conventional GWAS practice that accounts for population structure.

## Introduction

Genome-wide association studies (GWAS) aim to identify single nucleotide polymorphisms (SNPs) whose allelic variation is significantly tied to phenotypic variation. In principle, the tie between the allelic variation and phenotypic variation can be measured based on the variance among the phenotypic averages for all scores per each SNP (Kim, 2017; Kim, 2018a). Greater variance indicates a stronger tie. Conventional GWAS practice has been largely conducted using statistical methods such as the linear model and the linear mixed model (LMM). To date, the use of the LMM has been widely encouraged because of the general perception that accounting for a kinship matrix can reduce the noise between a phenotypic variable and an SNP variable, by correcting the bias that genetic relationship among entities in a population introduces (Yu et al, 2006; Bradbury et al, 2007; Kang et al, 2008; Lipka et al, 2012; Hoffman, 2013; Kim et al, 2018b). Recently, however, Kim (2019) demonstrated that the use of a kinship matrix actually makes the LMM unreliable. In this regard, this study excluded the LMM.

Conventional GWAS practice based on the linear model often regresses each SNP variable along with population-structure variables against a phenotypic variable, one by one across all SNPs. Therein, the use of population-structure variables aims to obtain an adjusted phenotype calculated by subtracting the estimated population-structure effect from the phenotype (Yu et al, 2006; Bradbury et al, 2007; Kang et al, 2008; Lipka et al, 2012; Hoffman, 2013; Kim et al, 2018b). For reliable GWAS practice, it is crucial to assure the adjusted phenotypes resulting across all SNPs are consistent. Otherwise, every SNP variable will be regressed against a differently adjusted phenotypic variable, which consequently confounds GWAS results. This study investigated whether accounting for population structure in the linear model for GWAS assures the adjusted phenotypes resulting across all SNPs to be consistent

## Materials and Methods

### Rice data set

This study used a rice data set comprising SNP data, principle component analysis (PCA) data and phenotypic data. The data set was originally used for GWAS by Zhao et al. (2011) and freely available to public at http://ricediversity.org/data/index.cfm. Therefore, more information about the data set can be found from the related paper. In the original data, 413 entities were genotyped with 36,901 SNPs. The number of SNPs was reduced to 12,983 by screening with a criterion of the minor allele frequency (MAF) of 0.1. The phenotype chosen for this study was seed length.

### Statistical model

The two linear models were established as follows:

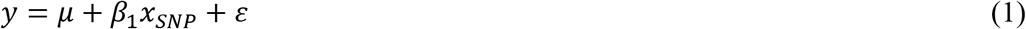

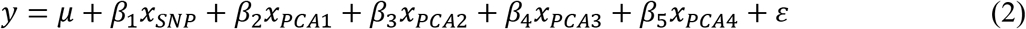

where *y* = the phenotypic observation; *μ* = the phenotypic mean; *x*_*SNP*_ = the SNP variable; *x*_*PCA1*_ = the PCA1 variable; *x*_*PCA2*_ = the PCA2 variable; *x*_*PCA3*_ = the PCA3 variable; *x*_*PCA4*_ = the PCA4 variable; *ε* = the error term; *β*_1_ = the coefficient for *x*_*SNP*_; *β*_2_ = the coefficient for *x*_*PCA1*_; *β*_3_ = the coefficient for *x*_*PCA2*_; *β*_4_ = the coefficient for *x*_*PCA3*_; *β*_5_ = the coefficient for *x*_*PCA4*_.

Equation 1 regresses the SNP variable against the phenotypic variable. Equation 2 regresses the SNP variable along with the four PCA variables (*x*_*PCA1*_, *x*_*PCA2*_, *x*_*PCA3*_, *x*_*PCA4*_) against the phenotypic variable. This means that Equation 2 regresses the SNP variable against the adjusted phenotypic variable obtained by accounting for the four PCA variables. Equation 3 highlights the adjusted phenotypic variable:

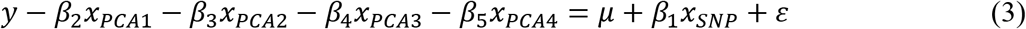

Equation 3 is compatible with Equation 2 and represents the adjusted phenotypic variable as *y* − *β*_2_*x*_*PCA1*_ − *β*_3_*x*_*PCA2*_ − *β*_4_*x*_*PCA3*_ − *β*_5_*x*_*PCA4*_.

### Manhattan plot

The F test was implemented as a significance test, from which P values were obtained. The P values transformed by −*log*_10_ were drawn in a Manhattan plot. It is important to note that the P values resulting from the linear model for GWAS are prone to genomic inflation. Prior to confirming the resultant Manhattan plot, therefore, it is necessary to calculate the genomic inflation factor (*λ*_*GC*_). The situation of *λ*_*GC*_ > 1 indicates the genomic inflation, which means that the resultant P values are overly estimated compared with the *χ*^2^-distribution (van Iterson et al, 2017). This study adjusted the genomic inflation using the genomic control. More information about the genomic control can be found in previous studies (Devlin and Roeder, 1999; Yang et al, 2011; van Iterson et al, 2017).

### Integrity validation of accounting for population structure in GWAS

Equation 3 (compatible with Equation 2) regresses each SNP variable against an adjusted phenotypic variable. As GWAS handle numerous SNPs one by one at a time, it is important to assure that the adjusted phenotypes resulting across all SNPs are consistent. Otherwise, each SNP variable will be regressed against a differently adjusted phenotypic variable. The consistency among the adjusted phenotypes resulting across all SNPs can be achieved, only if every coefficient per each PCA variable is consistent across all SNPs. To check the consistency among the adjusted phenotypes resulting across all SNPs, this study calculated Pearson coefficients between the phenotype and every adjusted phenotype.

### Data set and R code

All computations were conducted using R (R Core Team, 2016). The data set and R scripts used in this study are freely available at https://github.com/bongsongkim/Population.Structure.GWAS.

## Results

### Validation of consistency across all adjusted phenotypes

Table 1 summarizes the coefficients per each PCA variable, resulting from applying all SNPs to Equation 3. Figure 1 represents the estimated coefficients per each PCA variable, showing large variation. Figure 2 represents the estimated Pearson correlation coefficients between the phenotype and every adjusted phenotype, illustrating the adjusted phenotypes resulting across all SNPs are not consistent. This means that each SNP variable is regressed against a differently adjusted phenotypic variable.

**Table 1.** Summary of coefficients per each PCA variable in relation to Equation 3.

|  | Min. | 1<sup>st</sup> Qu. | Median | Mean | 3<sup>rd</sup> Qu. | Max |
| --- | --- | --- | --- | --- | --- | --- |
| $\beta_2$ | -7.833 | -2.246 | -2.162 | -2.109 | -2.024 | 3.629 |
| $\beta_3$ | -4.438 | -1.097 | -1.020 | -1.007 | -0.944 | 3.271 |
| $\beta_4$ | -13.610 | -9.229 | -9.194 | -9.180 | -9.160 | 0.659 |
| $\beta_5$ | -9.308 | 3.016 | 3.087 | 3.025 | 3.121 | 8.704 |

**Figure 1.**
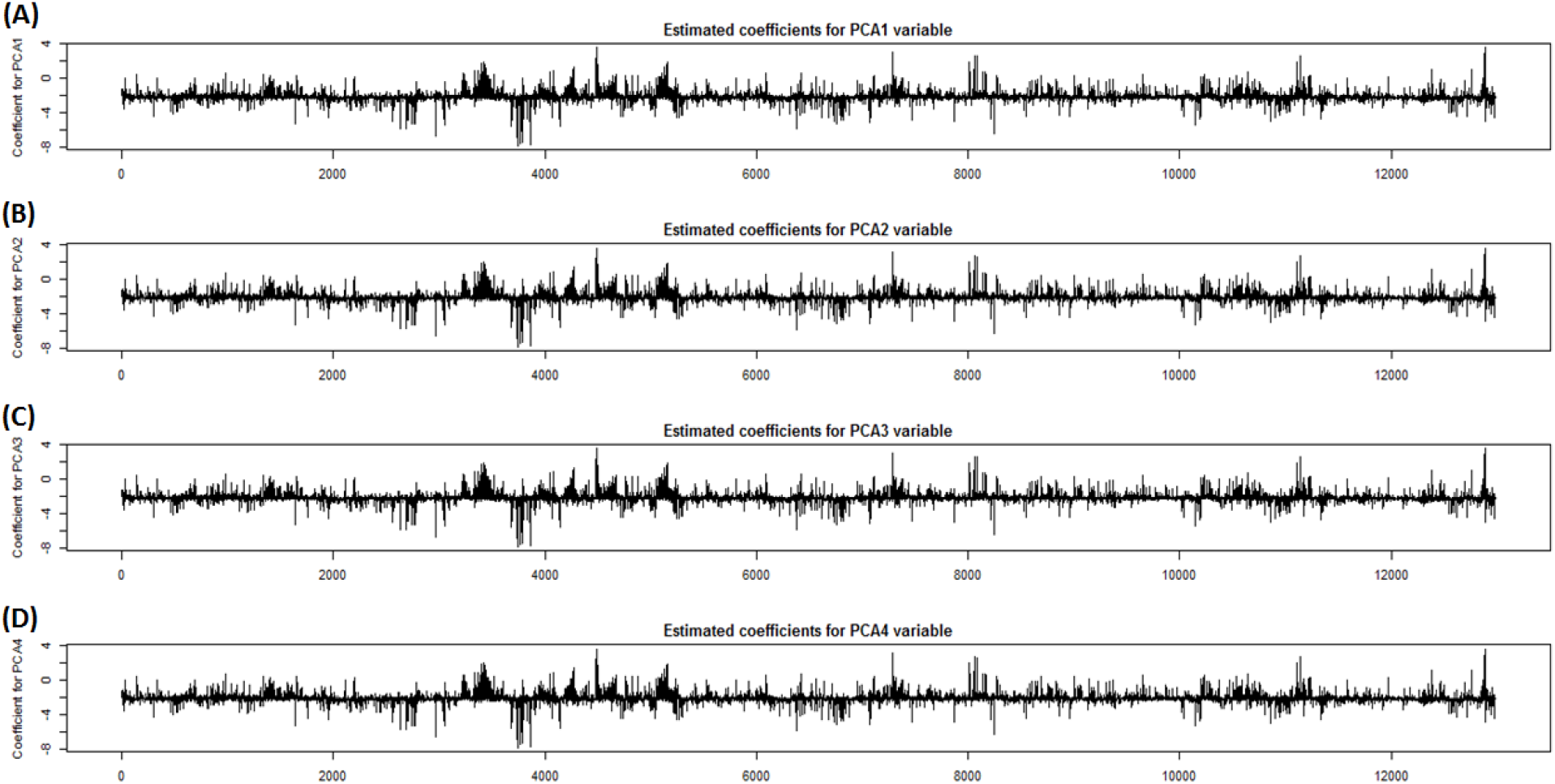
(A) Estimated coefficients for the PCA1 variable, (B) estimated coefficients for the PCA2 variable, (C) estimated coefficients for the PCA3 variable, (D) estimated coefficients for the PCA4 variable.

**Figure 2.**
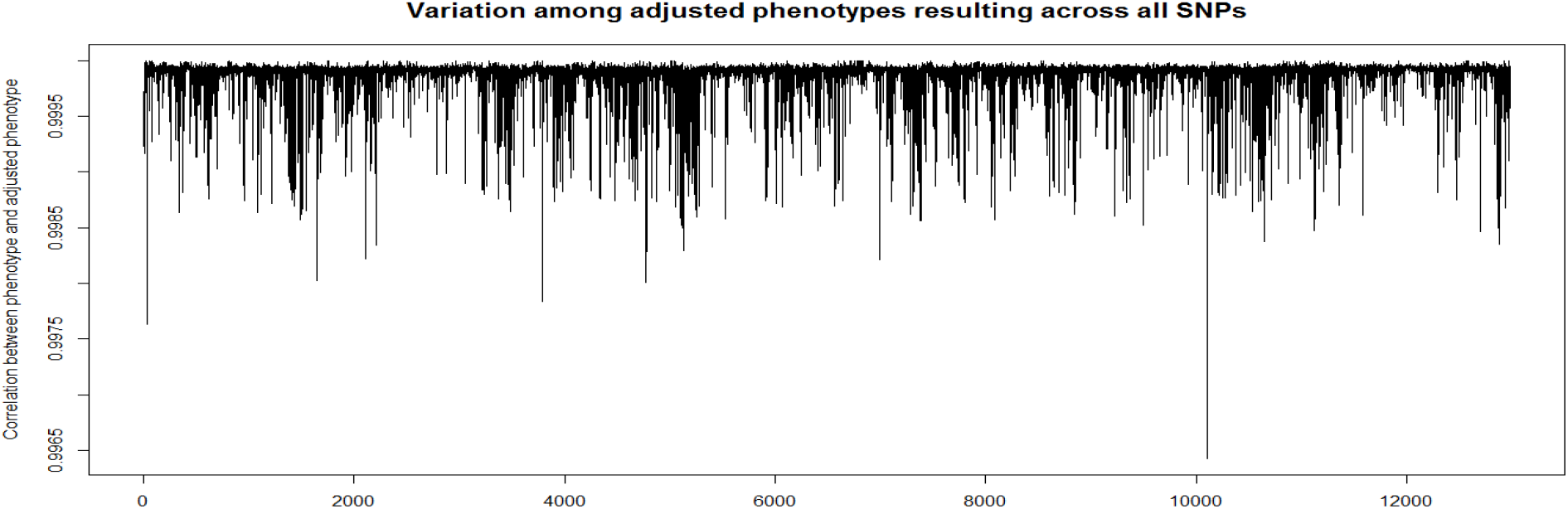
Pearson correlation coefficients between the phenotype and every adjusted phenotype.

### Impact of accounting for population structure in GWAS

Figure 3 shows four Manhattan plots, for which the same SNP and phenotypic data were used. Figure 3A represents the Manhattan plot in relation to Equation 1, in which the resultant λ_*GC*_ was 3.688. Figure 3C is the same as Figure 3A in shape. However, Figure 3C meets λ_*GC*_ = 1 by implementing the genomic control with Figure 3A. Figure 3E represents the Manhattan plot in relation to Equation 3, in which the resultant λ_*GC*_ was 1.433. Compared with Figure 3A, Figure 3E has substantially lower λ_*GC*_. This suggests that accounting for the four PCA variables was impactful in diminishing the genomic inflation. Figure 3G was obtained by adjusting Figure 3E by implementing the genomic control. This led to λ_*GC*_ = 1 in Figure 3G. It is apparent that Figure 3E has clearer background than Figure 3A in relation to accounting for the four PCA variables. In this regard, previous studies explained that accounting for population structure in the linear model for GWAS eliminates the noise in SNP-phenotype associations, which results in clear background in a resultant Manhattan plot (Yu et al, 2006; Kang et al, 2008; Korte and Farlow, 2013; Sul et al, 2018; Barton et al, 2019). However, Figure 4 illustrates that significant SNP-phenotype associations are not consistent between Figures 3C and 3G. This means that the clear background was not from eliminating the noise in SNP-phenotype associations, but from defining new SNP-phenotype associations.

**Figure 3.**
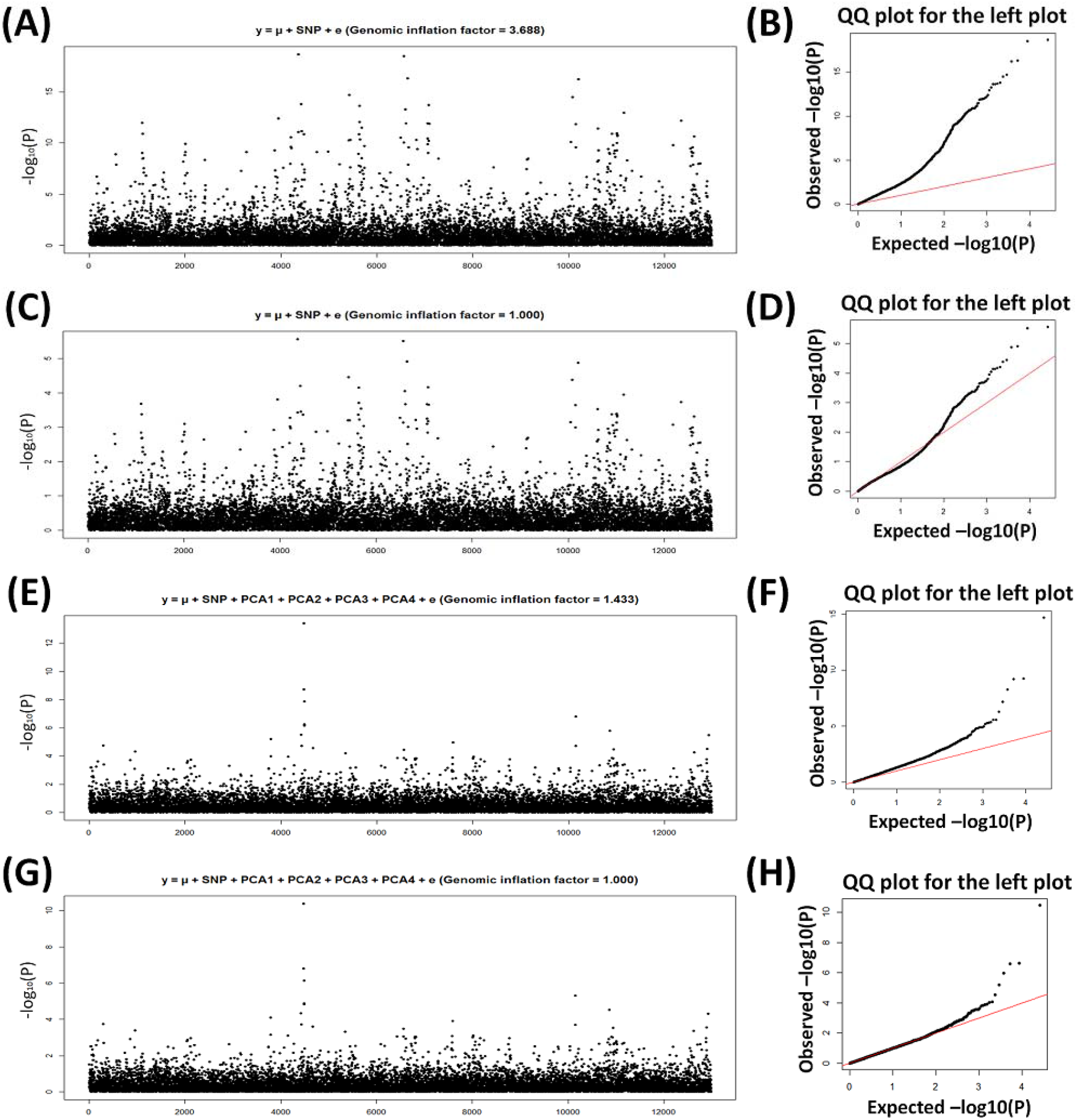
(A) Manhattan plot obtained by not accounting for the four PCA variables (*λ*_*GC*_ = 3.688), (C) Manhattan plot obtained by adjusting Figure 3A with implementing the genomic control (*λ*_*GC*_ = 1.000), (E) Manhattan plot obtained by accounting for the four PCA variables (*λ*_*GC*_ = 1.433), (G) Manhattan plot obtained by adjusting Figure 3C with implementing the genomic control (*λ*_*GC*_ = 1.000).

**Figure 4.**
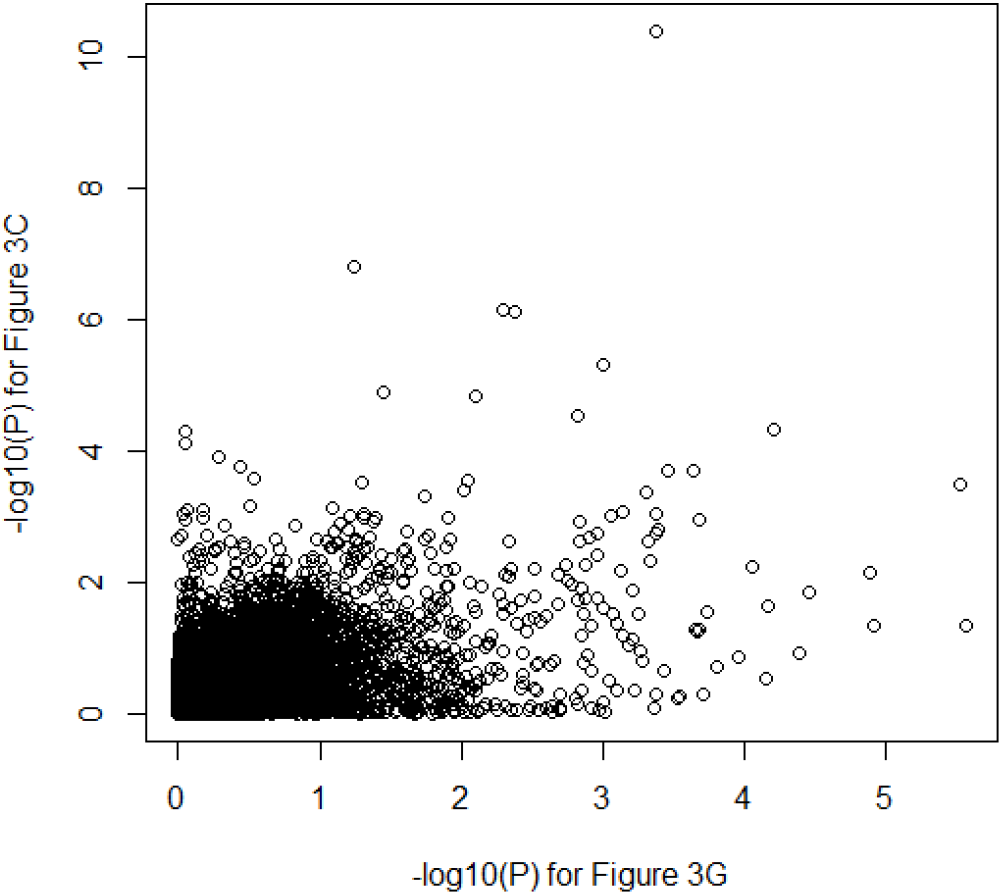
Correlation plot between the −log_10_ (P) values obtained by not accounting for the four PCA variables (Figure 3C) and the −log_10_ (P) values obtained by accounting for the four PCA variables (Figure 3G).

## Discussion

It is generally perceived that accounting for population structure in GWAS improves the quality of visual representation of a Manhattan plot by both suppressing genomic inflation and reducing false-positive SNP-phenotype associations (Yu et al, 2006; Bradbury et al, 2007; Kang et al, 2008; Lipka et al, 2012; Hoffman, 2013; Kim et al, 2018b). In fact, this study showed that accounting for the four PCA variables was very effective in diminishing the genomic inflation. Surprisingly, however, this study revealed that accounting for the four PCA variables breaks the consistency among the adjusted phenotypes resulting across all SNPs. The loss of the consistency consequently causes each SNP variable to be regressed against a differently adjusted variable, making the GWAS process unreliable. The use of population-structure variables in the linear model for GWAS implies two errors. First, the linear model is misused. Considering that the linear model is suited for analyzing data in experimental blocks, the use of continuous variables rather than factor variables necessarily causes an error. Second, the assumption for the relationship between phenotype and population structure is unjustified. The linear model for GWAS generally assumes that the population-structure variables additively contribute to the phenotypic variable. However, how the population structure biologically influences the phenotype has yet been unknown. Regardless of whether the additivity of the population-structure variables is true or false, the current way of accounting for population structure is inappropriate in that population-structure effects vary across all SNPs. The abovementioned errors consequently lead to the loss of the consistency among the adjusted phenotypes resulting across all SNPs and cause each SNP variable to be regressed against a differently adjusted phenotypic variable.

## Conclusion

The linear model assures to preserve the consistency among the adjusted phenotypes resulting across all SNPs, only if factor variables such as years, locations, replications and treatments are used. This study concluded that the conventional way of accounting for population structure makes the GWAS process unreliable. This is because the population structure is represented as continuous variables. If population structure can be represented as factor variables, accounting for the population structure in the linear model for GWAS will be sound.

